# Procaine and S-adenosyl-L-homocysteine (SAH) affect the expression of genes related to the epigenetic machinery and change the DNA methylation status of *in vitro* cultured bovine skin fibroblasts

**DOI:** 10.1101/574186

**Authors:** Schumann N.A.B., A.S. Mendonça, M.M. Silveira, L.N. Vargas, L.O. Leme, R.V. de Sousa, M.M. Franco

**Affiliations:** Institute of Genetics and Biochemistry, Federal University of Uberlândia, Uberlândia, Minas Gerais, Brazil; Laboratory of Animal Reproduction, Embrapa Genetic Resources and Biotechnology, Brasília, Distrito Federal, Brazil; School of Veterinary Medicine, Federal University of Uberlândia, Uberlândia, Minas Gerais, Brazil

**Author notes:** Email addresses: Naiara Araújo Borges Schumann, Anelise dos Santos Mendonça, Márcia Marques Silveira, Luna Nascimento Vargas, Ligiane Oliveira Leme, Regivaldo Vieira de Sousa.

## Abstract

Cloning using somatic cell nuclear transfer (SCNT) has many potential applications such as in transgenic and genomic-edited animal production. Abnormal epigenetic reprogramming of somatic cell nuclei is probably the major cause of the low efficiency associated with SCNT. Strategies to alter DNA reprogramming in donor cell nuclei may help improve the cloning efficiency. In the present study, we aimed to characterise the effects of procaine and S-adenosyl-L-homocysteine (SAH) as demethylating agents during the cell culture of bovine skin fibroblasts. We characterised the effects of procaine and SAH on the expression of genes related to the epigenetic machinery, including the DNMT1, DNMT3A, DNMT3B, TET1, TET2, TET3, and OCT4 genes, and on DNA methylation levels of bovine skin fibroblasts. We found that DNA methylation levels of satellite I were reduced by SAH (P=0.0495) and by the combination of SAH and procaine (P=0.0479) compared with that in the control group. Global DNA methylation levels were lower in cells that were cultivated with both compounds than in control cells [procaine (P=0.0116), SAH (P=0.0408), and both (P=0.0163)]. Regarding the transcriptional profile, there was a decrease in total DNMT transcript levels in cells cultivated with SAH and procaine. There was a higher level of TET3 transcripts in treated cells than in the controls. Our results showed that the use of procaine and SAH during bovine cell culture was able to alter the epigenetic profile of the cells. This approach may be a useful alternative strategy to improve the efficiency of reprogramming the somatic nuclei after fusion, which in turn will improve the SCNT efficiency.

## Introduction

Although two decades have passed since the first cloned mammal was born from an adult animal (1) and despite the somatic cell nuclear transfer (SCNT) technique having become a commercially available technique, its efficiency is still extremely low (2, 3). SCNT routinely involves the use of differentiated somatic cells, which display changed totipotency states through mechanisms dependent on epigenetic modifications (1, 4). Despite the successful cloning of several species, the use of differentiated somatic cells as donor nuclei is associated with a range of concerns, such as increased abortion rates, high embryonic lethality, and severe abnormalities in cloned foetuses and placentas in ruminants (5). Abnormal DNA methylation patterns represent one possible reason for the frequent developmental and metabolic anomalies and the very low survival rates of cloned calves (5–7).

Correct epigenetic reprogramming involves, among other modifications, the methylation of genomic DNA and post-translational modifications to histone proteins, which are involved in chromatin organisation (5, 8, 9). Epigenetic reprogramming during initial development is a dynamic process that involves two waves of methylation and demethylation and starts at the beginning of gametogenesis, continuing until the formation of all foetal tissues (9, 10). Some studies have reported that cloned embryos only partially demethylate their genomes and begin the *de novo* methylation process earlier than their counterparts (5, 7). Epigenetic processes such as DNA methylation and demethylation are regulated by different groups of enzymes. DNA methylation is performed by the DNA methyltransferase enzymes (DNMTs) (11). In fertilised embryos, through early development, methylation is reduced on or removed from numerous sequences, and from the 8-16 cell stage, a new embryonic methylation pattern generated by the *de novo* methyltransferases DNA methyltransferase 3 alpha (Dnmt3a) and DNA methyltransferase 3 beta (Dnmt3b) is established (5, 7). New patterns of DNA methylation on the genome are maintained by DNA methyltransferase 1 (DNMT1) and determine gene expression in embryo development (8). The mechanism of DNA demethylation involving the oxidisation of 5-methylcytosine (5mC) in 5-hydroxymethylcytosine (5hmC) is triggered by the ten-eleven translocation enzymes (TETs) (12). The TET family contains the TET1, TET2, and TET3 dioxygenases, which enable the conversion of 5-methylcytosine (5mC) to 5-hydroxylmethylcytosine (5-hmC) and 5-formylcytosine and 5-carboxylcytosine, ultimately leading to the replacement of the 5mC with a cytosine (13–15). TET proteins are also involved in histone modification, binding to metabolic enzymes and other proteins, influencing gene transcription (16). The difficulty in achieving efficient methylation reprogramming in SCNT embryos may be a consequence of the resistance of tissue-specific epigenetic patterns to be reprogrammed by the oocyte (17). Studies have reported that embryo development can be improved by reducing DNMT expression using epigenetic modifier agents and siRNA technology (18). It has also been found that SCNT embryo development can be significantly improved by strategies that stimulate changes in DNA methylation and histone modifications, such as the use of 5-Aza-2-deoxycytidine (zdC)(19–21), procaine (para-amino-benzoyl-diethylamino-ethanol) (22, 23), S-adenosyl-L-homocysteine (SAH) (24, 25), and Scriptaid (25), during cell culture. Procaine and SAH are demethylating compounds that are not analogous to nucleosides. Thus, they do not incorporate into the DNA molecule and therefore do not have cytotoxic effects, distinguishing them from most previously tested compounds (23, 26, 27). Procaine inhibits human cancer cell growth (27) through interactions with CpG-rich genomic regions and prevents the action of DNMT enzymes (23). Gao et al. (2009) showed that procaine can reverse the abnormal methylation of the Wif-1 gene and be used as a demethylating drug (28). SAH is a physiological by-product of the transmethylation reaction in cells involving S-adenosyl-L-methionine (SAM) and is found in the nucleus, cytoplasm, and extracellular environment (29). Increasing SAH concentrations promote its binding to the DNMT active sites, which in turn decreases DNA methylation (30). Jeon et al. (2008) demonstrated that SAH treatment of fibroblasts induces global DNA demethylation and increases the developmental potential for SCNT embryos with them also exhibiting higher telomerase activity levels, which is indicative of enhanced nuclear reprogramming (24).

Skin fibroblasts are the most commonly used donor cells used for nuclear transfer (NT) in cattle production. Thus, it is important to gain a better understanding of the dynamics of DNA methylation patterns in these cells in *in vitro* culture to improve SCNT protocols. Moreover, it is important to evaluate the dynamics of cells cultured with demethylating agents, as these culture conditions may offer promising prospects to verify the pluripotency status of treated cells and analyse the hypothesis that demethylating compounds could drive cells towards a dedifferentiated status.

Taken together, we aimed to evaluate the effects of procaine and/or SAH during the *in vitro* culture of bovine skin fibroblasts on the global and specific DNA methylation patterns and on the mRNA levels of the genes encoding the epigenetic machinery, such as DNMT1, DNMT3A, DNMT3B, TET1, TET2, TET3, and the pluripotency gene OCT4.

## Material and Methods

### Ethics statement

Experiments were performed in accordance with the Brazilian Law for Animal Protection and the Institutional Guidelines for Animal Care and Experimentation. The project was approved by the Ethics Committee in Animal Use at EMBRAPA Genetic Resources and Biotechnology (CEUA/CENARGEN) at a special meeting held on July 2, 2015.

### Experimental design

Adult skin fibroblasts were obtained from a biopsy taken from a male Nellore (*Bos taurus indicus*) bull. Procaine and SAH were used alone or in combination for 2 weeks during bovine skin fibroblast *in vitro* culture in four experimental treatments. Four biological replicates were performed for each treatment: (1) control, (2) cells cultured with 1.0 mM procaine, (3) cells cultured with 1.0 mM SAH, and (4) cells cultured with 1.0 mM SAH and 1.0 mM procaine. We specifically focused on evaluating the effects of procaine and SAH on cell growth, specific and global DNA methylation, and the transcript levels of genes encoding the epigenetic machinery, including DNMT1, DNMT3A, DNMT3B, TET1, TET2, and TET3. In addition, to verify the pluripotency status of the treated cells, we analysed the expression of the OCT4 gene. The experimental design is presented in Fig. 1.

**Fig. 1.**
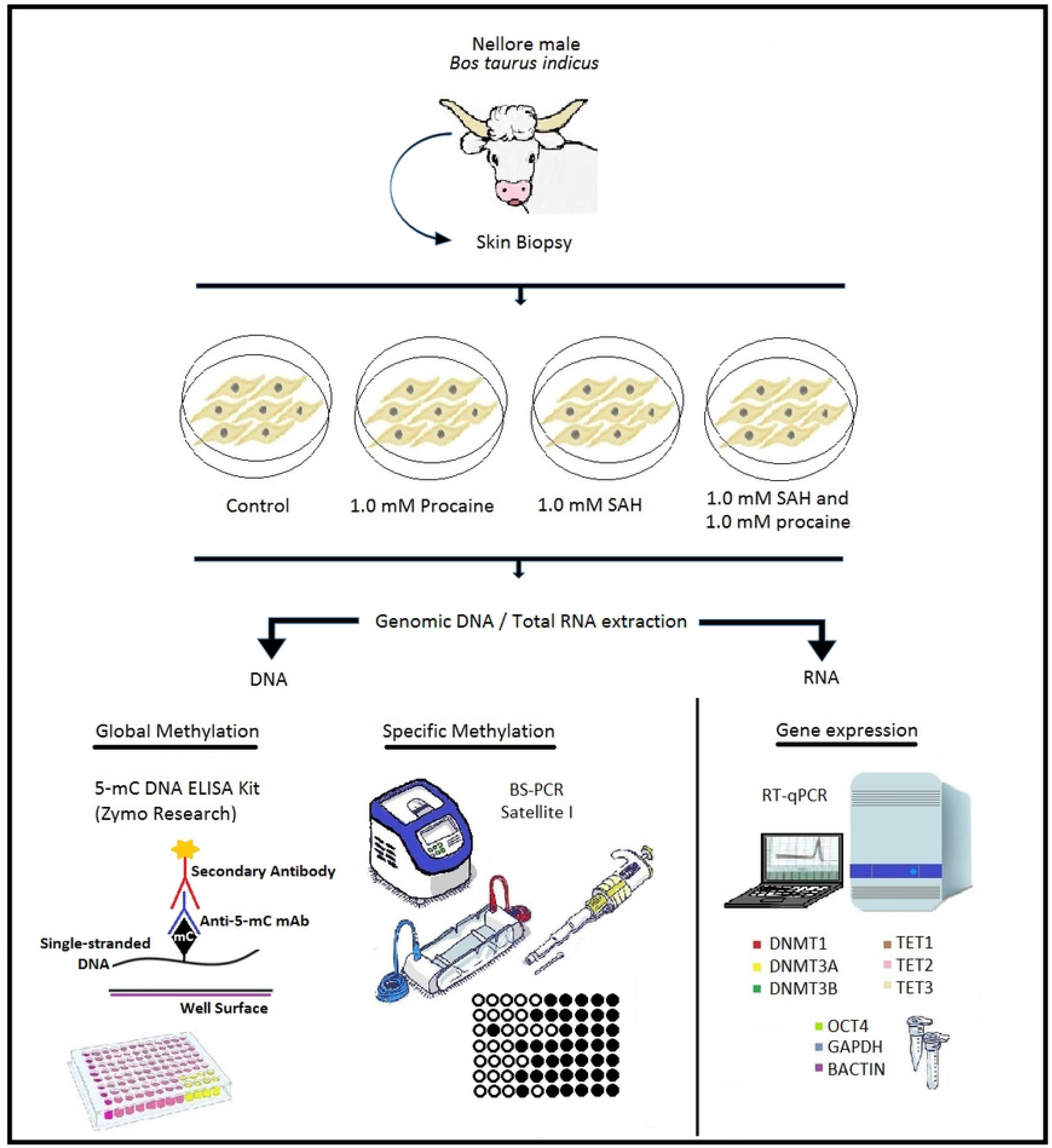
Scheme of the experimental design. Skin biopsy was obtained from a male Nellore (*Bos taurus indicus*) bull. From this biopsy, an *in vitro* culture of skin fibroblasts was established. Cells from the control group were cultured only with DMEM. The other groups were supplemented with procaine, S-adenosyl-L-homocysteine (SAH), or a combination of both substances for 2 weeks. After 2 weeks of *in vitro* culture, genomic DNA and total RNA extractions were performed. Genomic DNA was used for global (5-mC DNA ELISA Kit -Zymo Research) and specific (satellite I region – BS-PCR) methylation analyses. Total RNA was used for analysis of gene expression by RT-qPCR. Genes involved in the epigenetic machinery (DNMT1, DNMT3A, DNMT3B, TET1, TET2, and TET3) and pluripotency status (OCT4) were analysed. The β-actin and GAPDH genes were used as endogenous controls.

### Cell culture establishment

The skin biopsy was cut into small pieces (2–3 mm^2^), and the explants were cultured in Dulbecco’s modified Eagle Medium (DMEM-Gibco^®^ Life Technologies, Carlsbad, CA, USA) supplemented with 3.7 g/L sodium bicarbonate, 110 mg/L pyruvate, 10% foetal bovine serum (FCS), and antibiotics in 25 cm2 bottles at 39°C in a humidified atmosphere with 5% CO2 until confluence. After establishing the cell culture, cells were frozen (second passage) in DMEM containing 10% dimethyl sulphoxide (DMSO, Gibco, Carlsbad, CA, USA). Cell pallets were stored at −80°C for 24 h and subsequently transferred to liquid nitrogen, where they were stored until use.

### Cell culture using procaine and/or SAH

Cells were *in vitro* cultured using procaine or SAH according to a previously established protocol in the Laboratory of Animal Reproduction (Embrapa Genetic Resources and Biotechnology, Brazil). Briefly, cells were thawed, subcultured, and grown until they reached confluence in DMEM supplemented with 3.7 g/L sodium bicarbonate, 110 mg/L pyruvate, 10% foetal bovine serum (FBS), and antibiotics for 2 weeks. The culture media also contained 1.0 mM procaine, 1.0 mM SAH, or a combination of the two compounds, as described in the experimental design (Fig. 1). Cells were evaluated after reaching confluence using an inverted Axiovert 135M microscope at 10X magnification (ph1) with an attached CFI60 Nikon optical system and an ECO-LED lighting system.

### DNA isolation

*In vitro* cultured cells were used for genomic DNA isolation for use in methylation analyses. Genomic DNA was isolated using the DNeasy^®^ Blood & Tissue Kit (Qiagen, Hilden, Germany) according to manufacturer’s protocol. DNA samples were diluted in 20 μL buffer AT and stored at −20°C after measuring the DNA concentration.

### Sodium bisulphite treatment

Genomic DNA was treated with sodium bisulphite using the EZ DNA Methylation-Lightning™ Kit (Zymo Research, Orange, CA, USA) according to manufacturer’s protocol. After bisulphite treatment, the DNA samples were stored at −80°C until PCR amplification or global methylation analyses.

### PCR of bisulphite-converted DNA

Bisulphite-converted DNA samples were submitted to PCR amplification. The primers were designed with MethPrimer software to flank and amplify a CpG island in the repetitive DNA sequence of the *Bos taurus* bovine testis satellite I (satellite I)(31). PCR was performed in a total volume of 20 μL using 1x Taq buffer, 1.5 mM MgCl2, 0.4 mM dNTPs, 1 U Platinum™ Taq polymerase (Invitrogen, CA, USA), 0.5 μM each primer (forward and reverse), and 2 μL of bisulphite-treated DNA. The following temperature and time conditions were used: (1) an initial denaturing step at 94°C for 3 min, (2) 40 cycles at 94°C for 40 s, 45°C for 1 min, and 72°C for 1 min, and (3) a final extension at 72°C for 15 min.

### Cloning and bisulphite sequencing

After PCR, the amplicons were purified from an agarose gel using the Wizard SV Genomic DNA Purification System (Promega Corp., Madison, WI, USA) according to the manufacturer’s protocol. Then, the purified amplicons were inserted into the TOPO TA Cloning vector (pCR™II-TOPO^®^ vector system, Invitrogen, Carlsbad, CA, USA) and transferred into DH5α cells using a heat shock protocol. Then, plasmid DNA was isolated using the QIAprep Spin Miniprep Kit (Qiagen, CA, USA), and individual clones were sequenced using BigDye^®^ cycle sequencing chemistry.

The sequencing quality was analysed using Chromas^®^, and the methylation patterns were analysed using the BiQ Analyser program (32) (MPI for Informatics, Saarland, Germany). The DNA sequences were compared with a reference sequence from GenBank, accession number AH001157.2. Only sequences that originated from clones with ≥ 95% homology and cytosine conversion were used.

### Global DNA methylation analysis

Genomic DNA samples were also used for global DNA methylation analysis using the 5-mC DNA ELISA Kit (Zymo Research, Irvine, CA, USA) according to manufacturer’s protocol. Briefly, each DNA sample (∼100 ng), in triplicate, was bound to strip-wells that were specifically treated to have high DNA affinity. Global DNA methylation was detected using specific antibodies and was quantified colorimetrically by reading the absorbance at 450 nm in a microplate spectrophotometer (Bio-Rad Microplate Reader, Bio-Rad Laboratories, Redmond, WA, USA). The percentage of methylated DNA was proportional to the measured optical density (OD) intensity. Relative quantification was used to calculate the percentage of 5-mC in the total cytosine content in the bovine genome.

### Total RNA isolation and cDNA synthesis

Total RNA isolation was performed in biological quadruplicates of exponentially growing fibroblasts using the PureLinkTM RNA Mini kit (Invitrogen, USA) following the manufacturer's protocol. Total RNA was eluted in 20 μL of DEPC-treated water. Immediately before use for cDNA synthesis, total RNA was treated with 2 U of RQ1 RNase-Free DNase (Promega, USA) at 37°C for 30 min, and the DNase was inactivated by incubation at 65°C for 10 min. cDNA was synthesised using 1 µg of total RNA in a 20 µL final volume using the GoScript Reverse Transcription System (Promega, USA) according to the manufacturer’s protocol. cDNA synthesis was performed in a Veriti 96-Well Thermal Cycler Gradient (Applied Biosystems) using the following conditions: 5 min at 25°C, 60 min at 42°C, 15 min at 70°C, and holding at 4°C. cDNA samples were stored at −20°C until use.

### Real Time PCR

RT-qPCR reactions were performed in triplicate in a final volume of 25 µL containing 1 µL of cDNA, 12.5 µL of GoTaq^®^ qPCR Master Mix (Promega, USA), and 0.2 µM each primer in a 7500 Fast Real-Time PCR System (Applied Biosystems, USA). RT-qPCR conditions were as follows: 95°C for 20 s and 40 cycles of a denaturation step at 95°C for 3 s and an annealing/extension step at 60°C for 30 s. Reactions were optimised to provide a maximum amplification efficiency for each gene using a relative standard curve, and the efficiencies of each pair of primers are shown in Table 1. Each sample was analysed in triplicate, and the specificity of each PCR product was determined by analysing the melting curve and the size of amplicons in an agarose gel. The glyceraldehyde-3-phosphate dehydrogenase (GAPDH) and β-actin (BACT) genes were used for data normalisation using the geometric means of their Ct (cycle threshold) and their efficiencies of amplification. The control group (cells cultivated without procaine and SAH) was used as the reference sample. The relative abundance of mRNA for each gene was calculated and compared among the experimental groups using the ∆∆Ct method, with efficiency correction using the Pfaffl method (33).

**Table 1:**
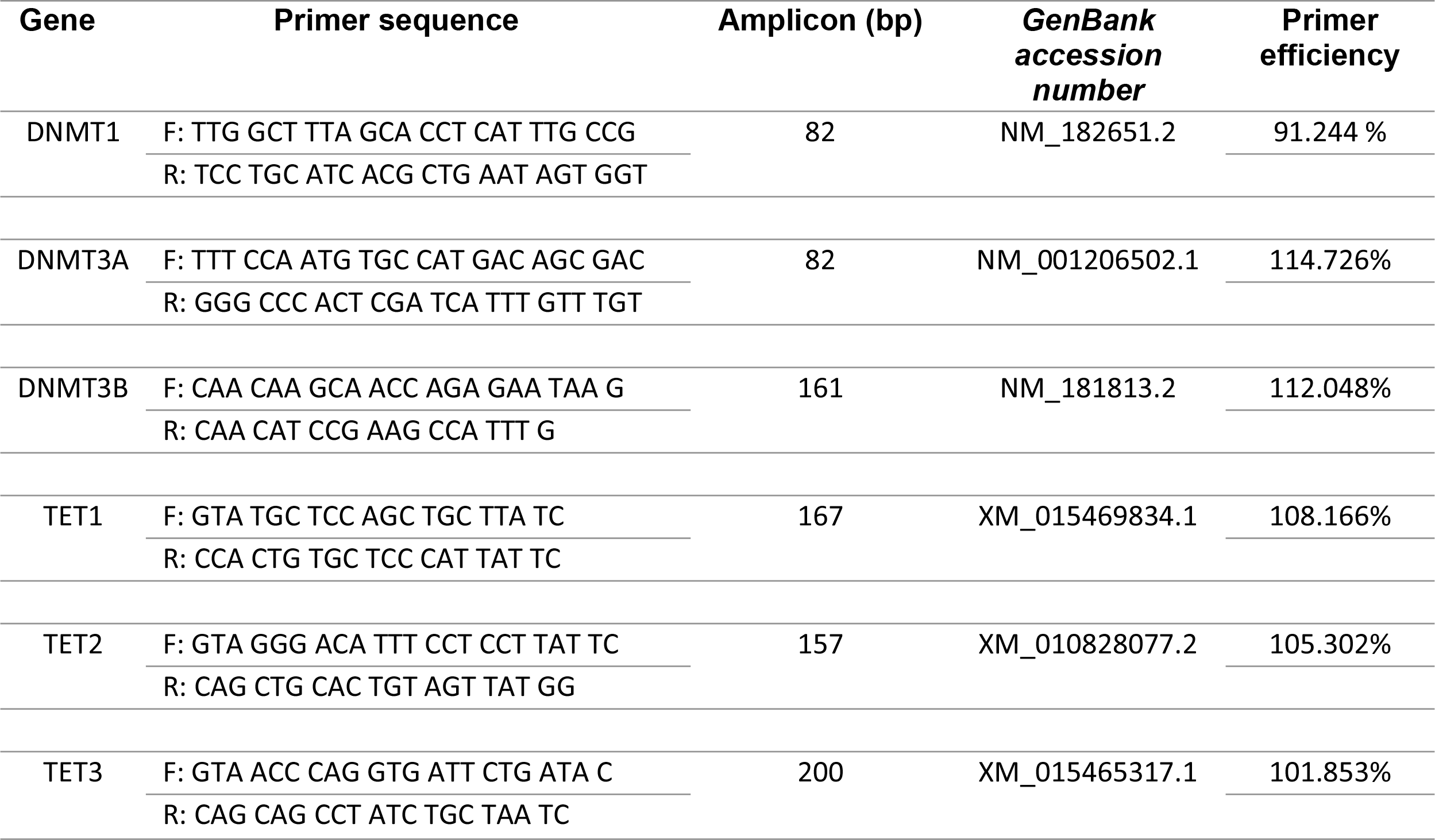
Primers for gene expression analysis

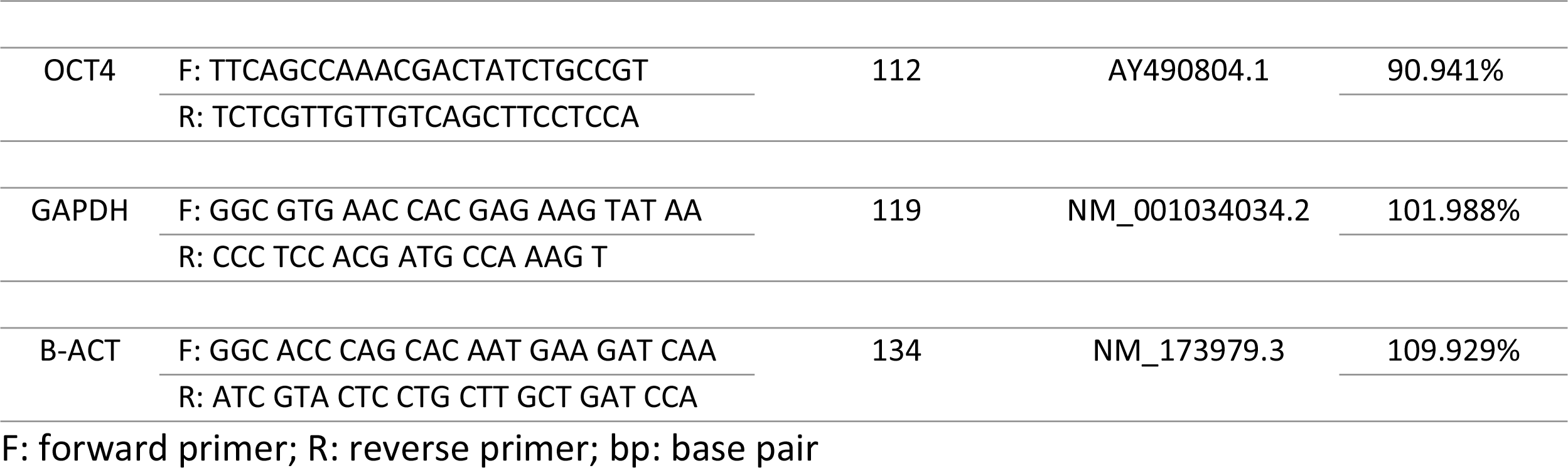

Six genes related to DNA methylation reprogramming were selected for analysis, DNMT1, DNMT3A, DNMT3B, TET1, TET2 and TET3. In addition, OCT4, related to pluripotency status, was also analysed. Details on the genes and primers are presented in Table 1.

### Statistical analyses

All analyses were performed using GraphPad Prism software (Version 6.0). Data were compared among experimental groups using ANOVA and Tukey’s test or the Kruskal-Wallis and Mann-Whitney tests for data showing or not showing normality, respectively. The normality of the data was analysed using the Shapiro-Wilk test. The results are presented as the mean ± the standard deviation.

## Results

### Cell culture and morphology

After *in vitro* culture, the cells from all treatments showed normal cell growth and reached confluence. No morphological changes were observed during culture (Fig. 2).

**Fig. 2.**
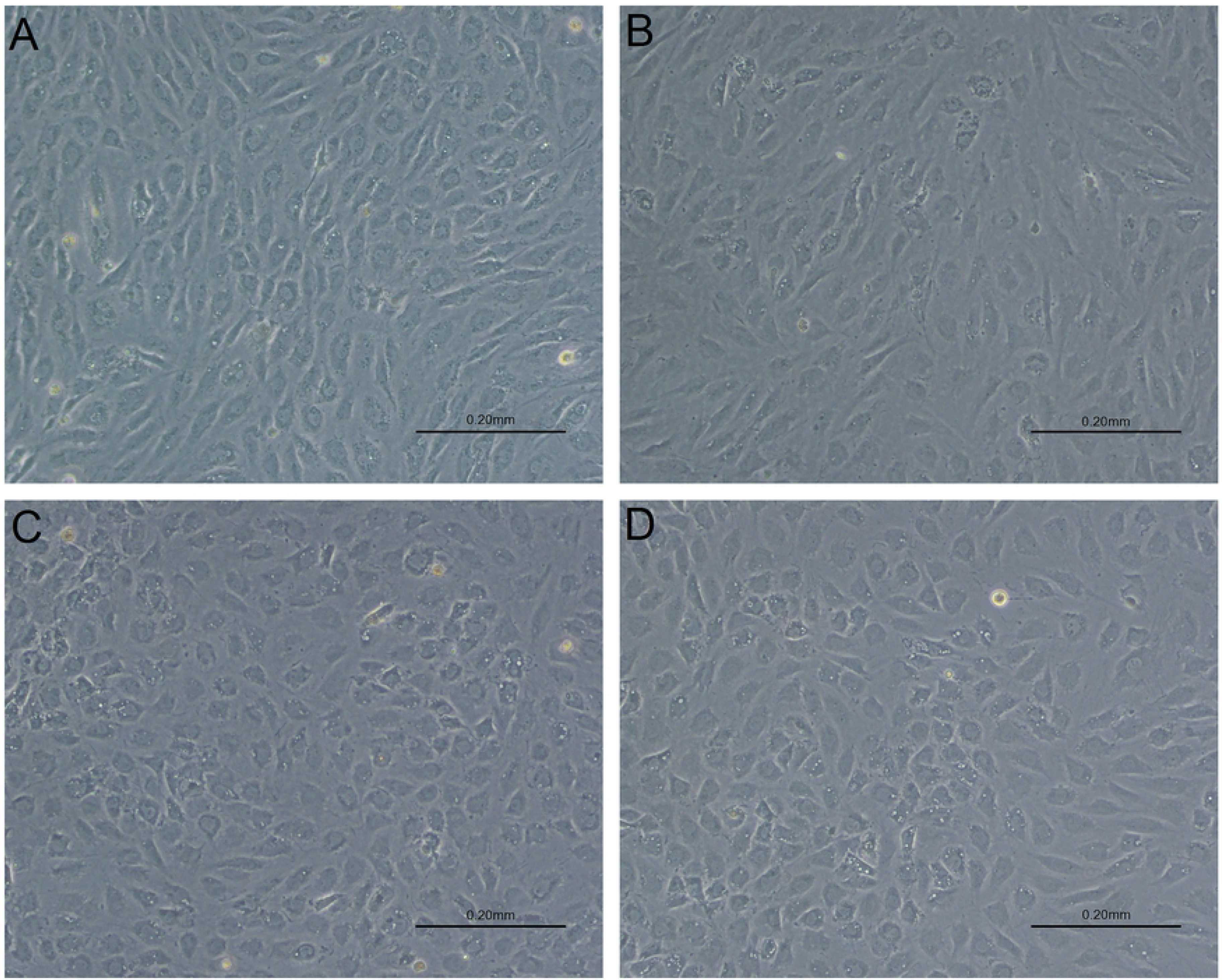
Photomicrographs of bovine skin fibroblast cell lines in *in vitro* culture. A - Control group. B - Culture supplemented with 1.0 mM procaine. C – Culture supplemented with 1.0 mM S-adenosyl L-homocysteine (SAH). D - Culture supplemented with 1.0 mM procaine and 1.0 mM SAH. 100x magnification. Bar: 0.20 mm.

### Total cell counting

The cells were separately counted after treatment with procaine, SAH, or procaine and SAH. The total cell number observed for each treatment, in quadruplicate, is shown in Table 2 and Fig. 3. No differences were found among the treatments (P=0.0899).

**Table 2:**
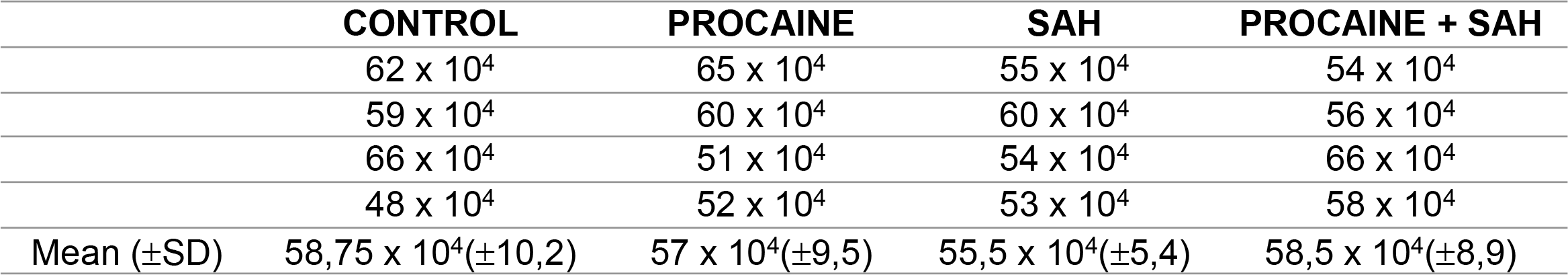
Total number of cells from the *in vitro* culture of bovine skin fibroblasts cultured with procaine, SAH, or procaine and SAH counted in a Neubauer chamber.

**Fig. 3.**
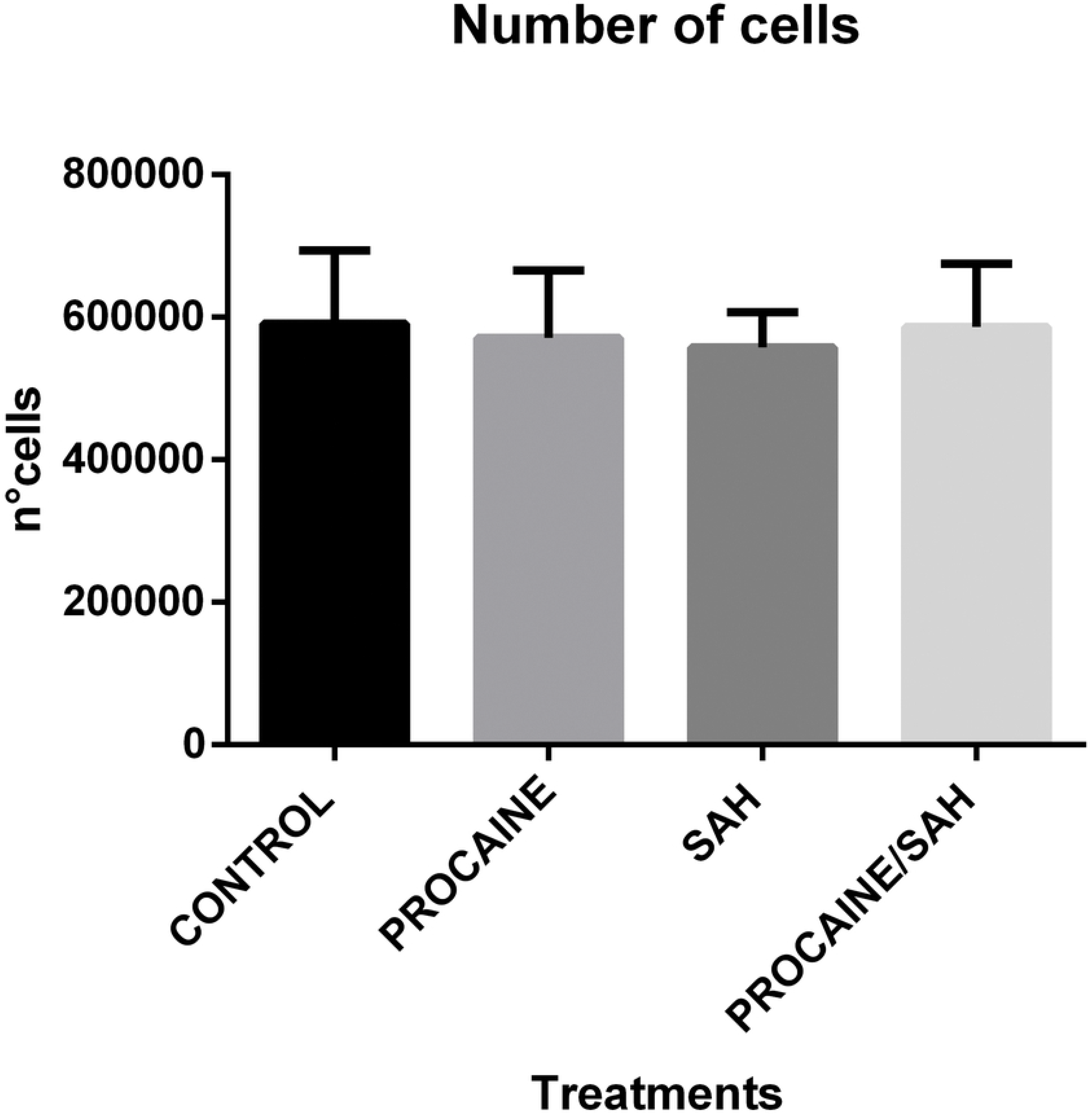
Mean ± standard deviation (four biological replicates) of the number of cells for each treatment (procaine, S-adenosyl L-homocysteine (SAH), procaine and SAH, and control groups). The cells were counted in a Neubauer chamber.

### DNA methylation profile of satellite I

The bisulphite sequencing results are shown in Fig. 4 and Fig. 5. The DNA methylation status of satellite I, a DNA repeat element, was investigated. We found that the genomic DNA from cells cultured for 2 weeks with SAH and with procaine + SAH was less methylated compared to that from cells in the control group (P=0.0495 and P=0.0479, respectively; Fig. 4 and Fig. 5). However, no differences were found for cells cultured with procaine alone.

**Fig. 4.**
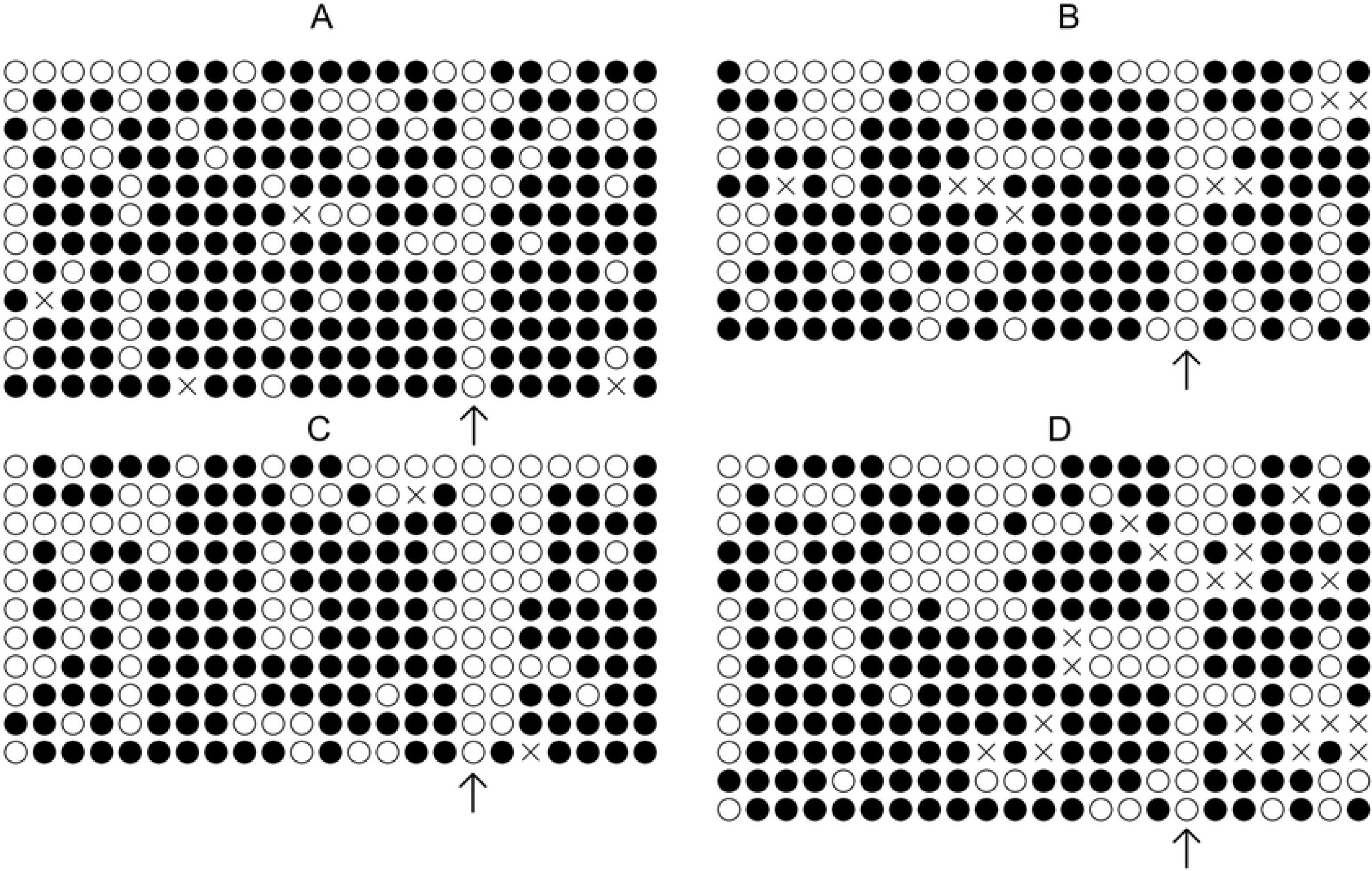
DNA methylation pattern of the satellite I region in bovine skin fibroblasts. A - Control cells. B - Cells cultured with 1.0 mM procaine. C - Cells cultured with 1.0 mM S-adenosyl L-homocysteine (SAH). D - Cells cultured with 1.0 mM procaine and SAH. Each line represents one individual clone, and each circle represents one CpG dinucleotide (a total of 23 CpGs were analysed). White circles represent unmethylated CpGs, filled black circles represent methylated CpGs, and grey circles represent CpGs that could not be analysed. Arrows indicate that for all treatments, the cytosines in position 17 were always demethylated.

**Fig. 5.**
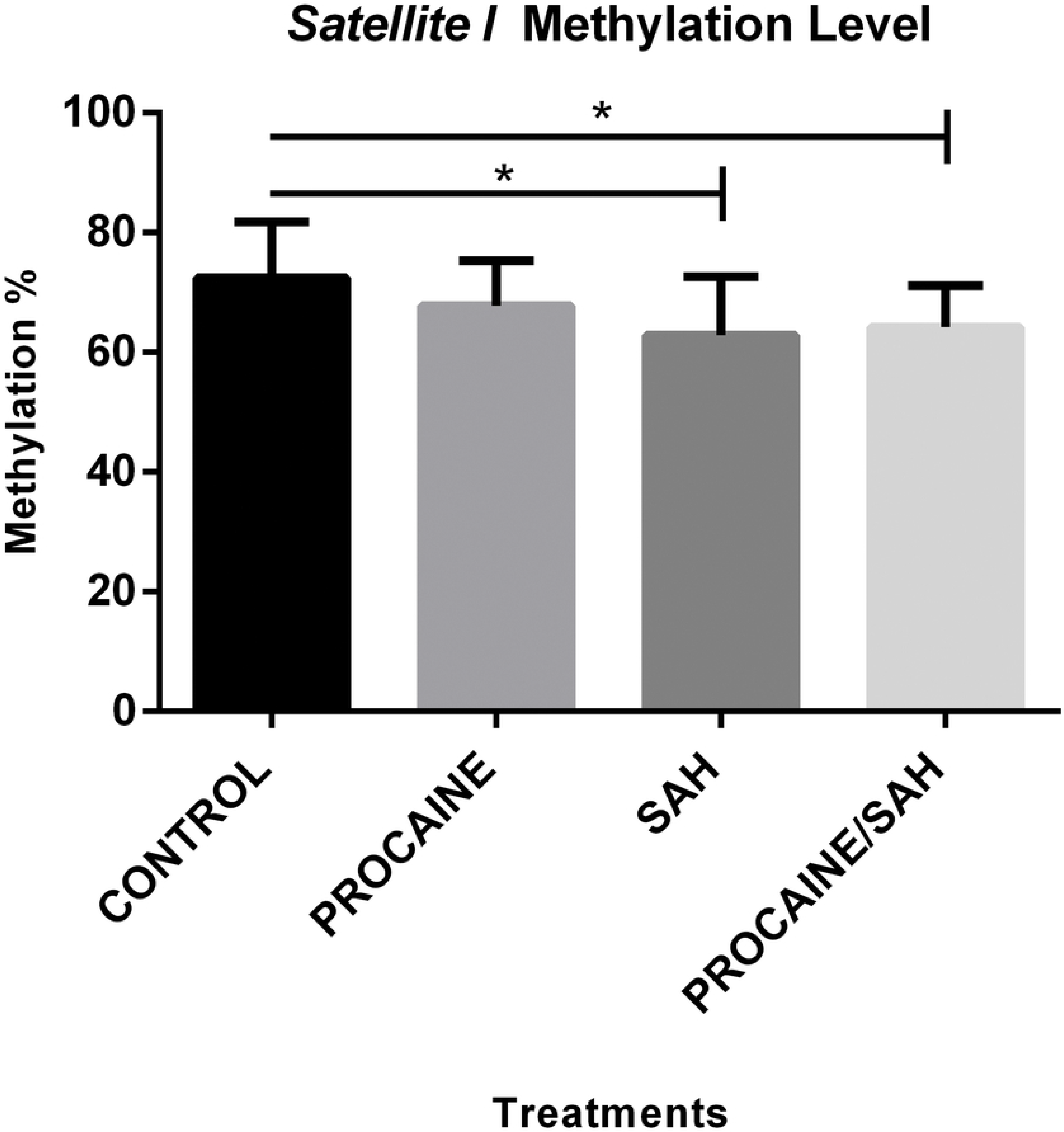
Mean ± standard deviation of DNA methylation levels of the satellite I region in bovine skin fibroblasts. Control cells and cells cultured with procaine, S-adenosyl L-homocysteine (SAH), or procaine and SAH. An asterisk represents a significant difference.

### Global methylation analysis

We found that global methylation levels were lower in all treated cell groups, procaine (P=0.0116), SAH (P=0.0408), and procaine and SAH (P=0.0163), than in the control group (Fig. 6).

**Fig. 6.**
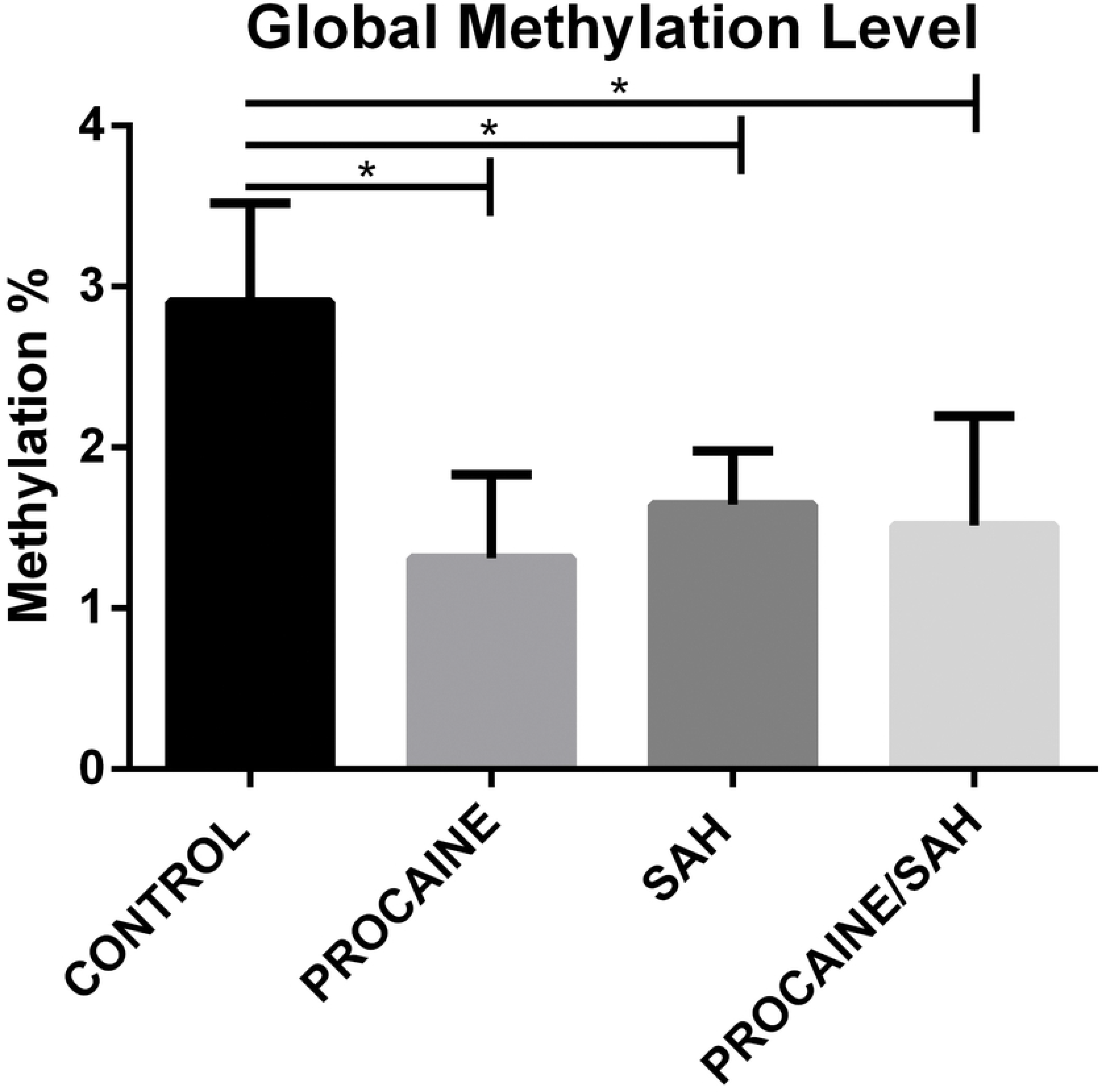
Mean ± standard deviation of global DNA methylation levels in bovine skin fibroblasts cultured with procaine, S-adenosyl L-homocysteine (SAH), or procaine and SAH. Biological quadruplicates and technical triplicates were performed for each measurement.

### Quantification of relative mRNA abundance

We quantified the mRNA levels of genes involved in the epigenetic methylation process, DNMT1, DNMT3A, DNMT3B, TET1, TET2, and TET3, and pluripotency status, OCT4. The mRNA levels of all genes that were evaluated were detected in bovine skin fibroblasts. Specific amplicon sizes determined in agarose gels and specific melting temperatures for each gene studied confirmed the high specificity of the primers that were used in this study (Fig. 7 and Fig. 8). When analysing each DNMT individually, per each treatment, no differences were found among the groups, but when considering the expression of each DNMT individually, regardless of the treatment, the mRNA levels of DNMT1, DNMT3A, and DNMT3B were lower in cells cultivated with both substances (Fig. 9). Moreover, when considering the expression of the group of three DNMTs and analysing per treatment individually, procaine, SAH, and procaine in combination with SAH decreased the mRNA levels of DNMTs (Fig. 10). Regarding the TET genes, the only significant difference that was verified was for TET3, for which there were higher levels of transcripts in cells cultivated with both substances, regardless of the treatment, when compared to the control (Fig. 11). Figure 12 shows the log-fold change for all analysed genes related to DNA methylation reprogramming, presenting the downregulated or upregulated profile of each gene compared to that of the control (Fig. 12). For the OCT4 gene, we verified that there was no significant difference in treated cells in relation to the control group (Fig. 13 and Fig. 14).

**Fig. 7.**
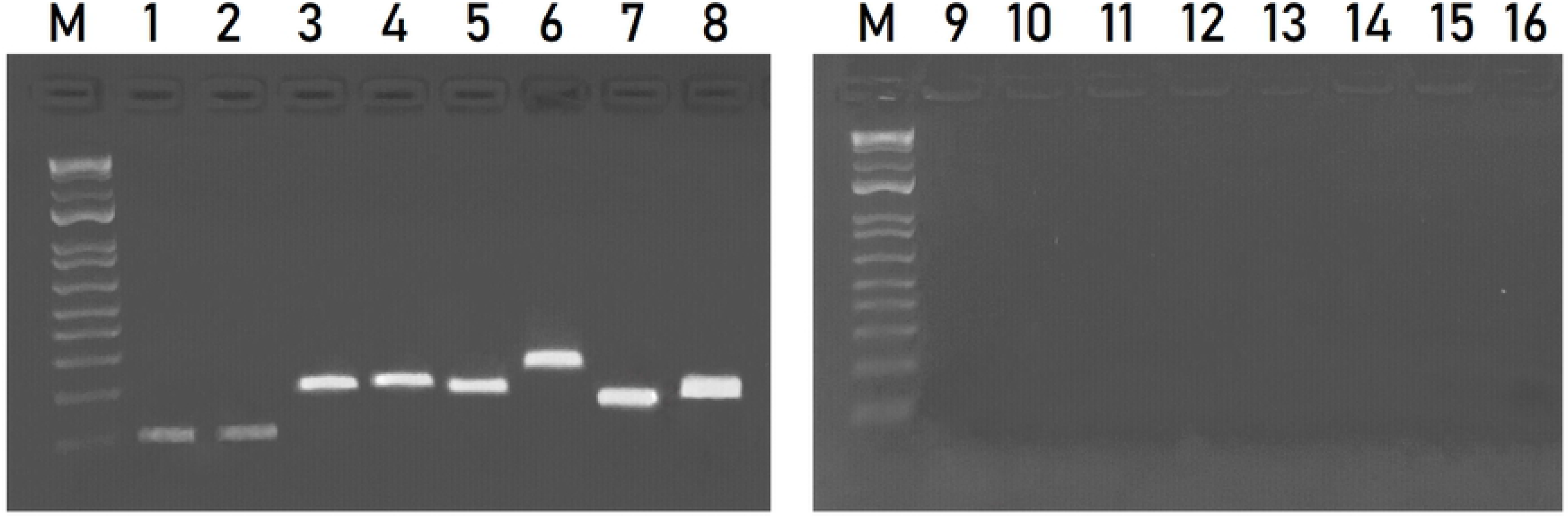
Agarose gel (1.5%) showing amplicons relative to the genes evaluated. Lane 1 - DNMT1 (82 bp); lane 2 - DNMT3A (82 bp); lane 3 - DNMT3B (161pb); lane 4 - TET1 (167 bp); lane 5 - TET2 (157 bp); lane 6 - TET3 (200 bp); lane 7 - GAPDH (119 bp); and lane 8 - β-ACTIN (134 bp). M: 100 bp DNA ladder. Lanes 9 to 16 represent the negative control PCR for each gene, respectively. bp – base pair.

**Fig. 8.**
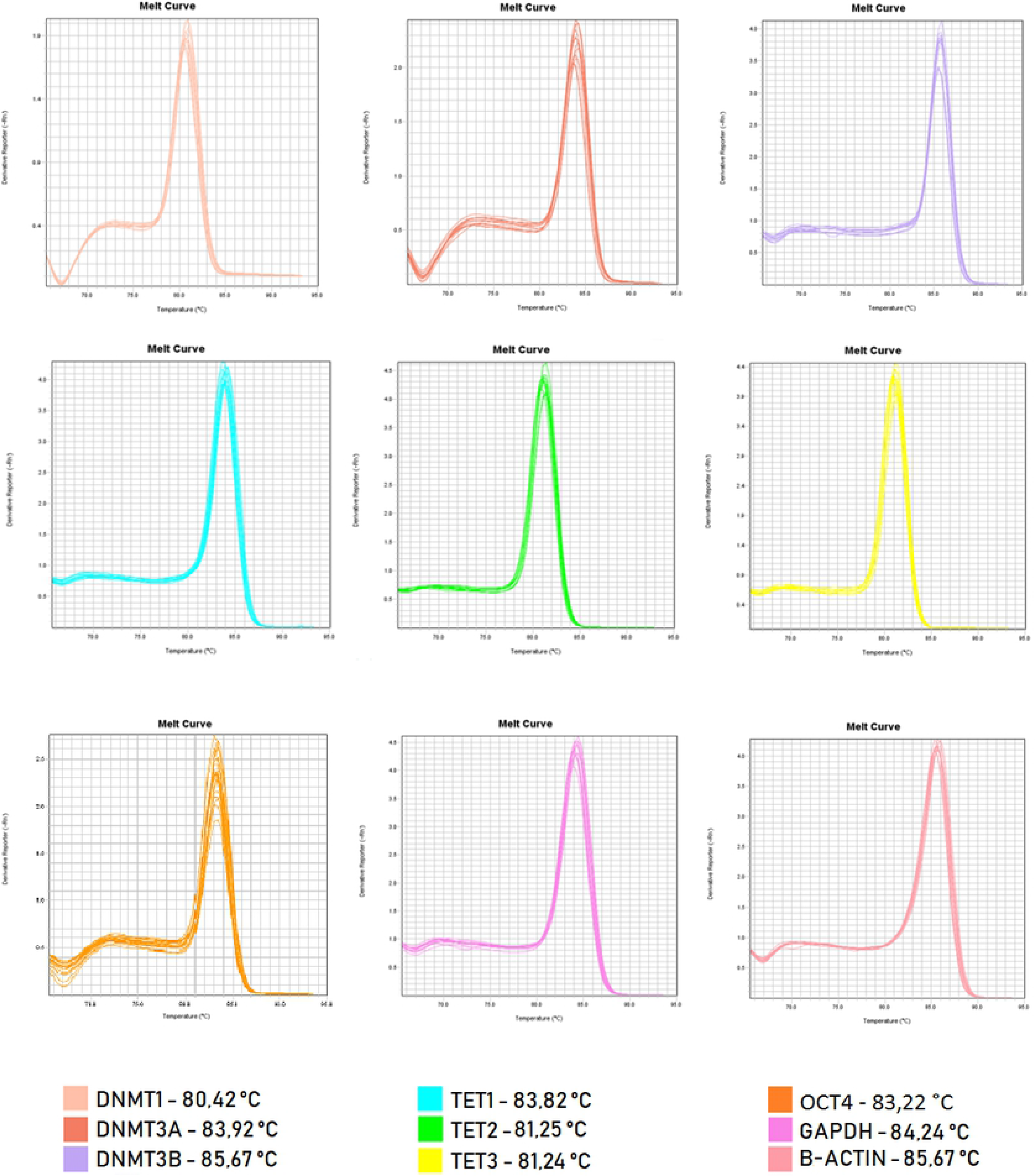
Melting curves for primers used in RT-qPCR.

**Fig. 9.**
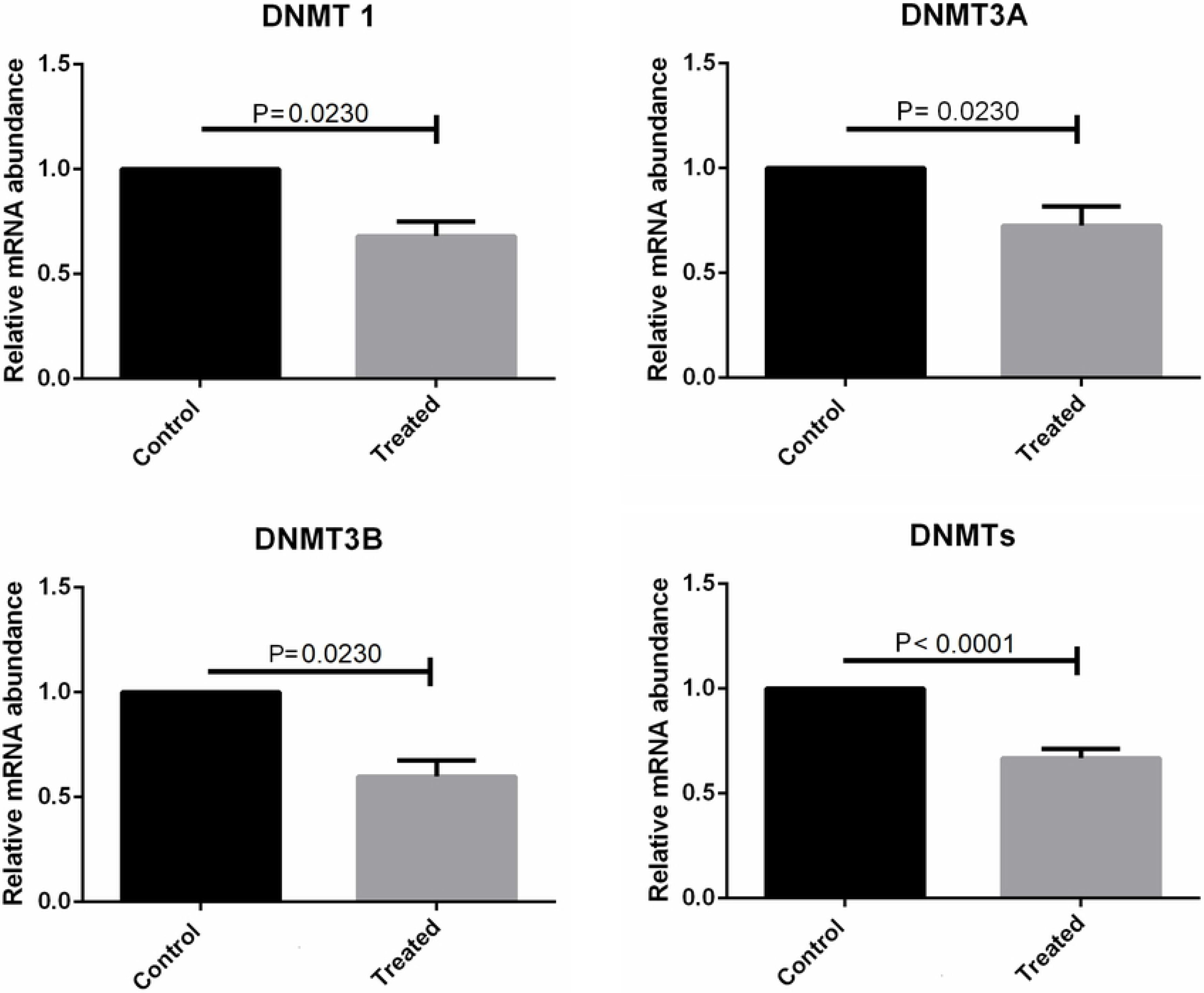
mRNA levels of DNMT1, DNMT3A, and DNMT3B determined by RT-qPCR in bovine skin fibroblasts treated with procaine and/or S-adenosyl L-homocysteine (SAH). Differences were considered significant when p <0.05.

**Fig. 10.**
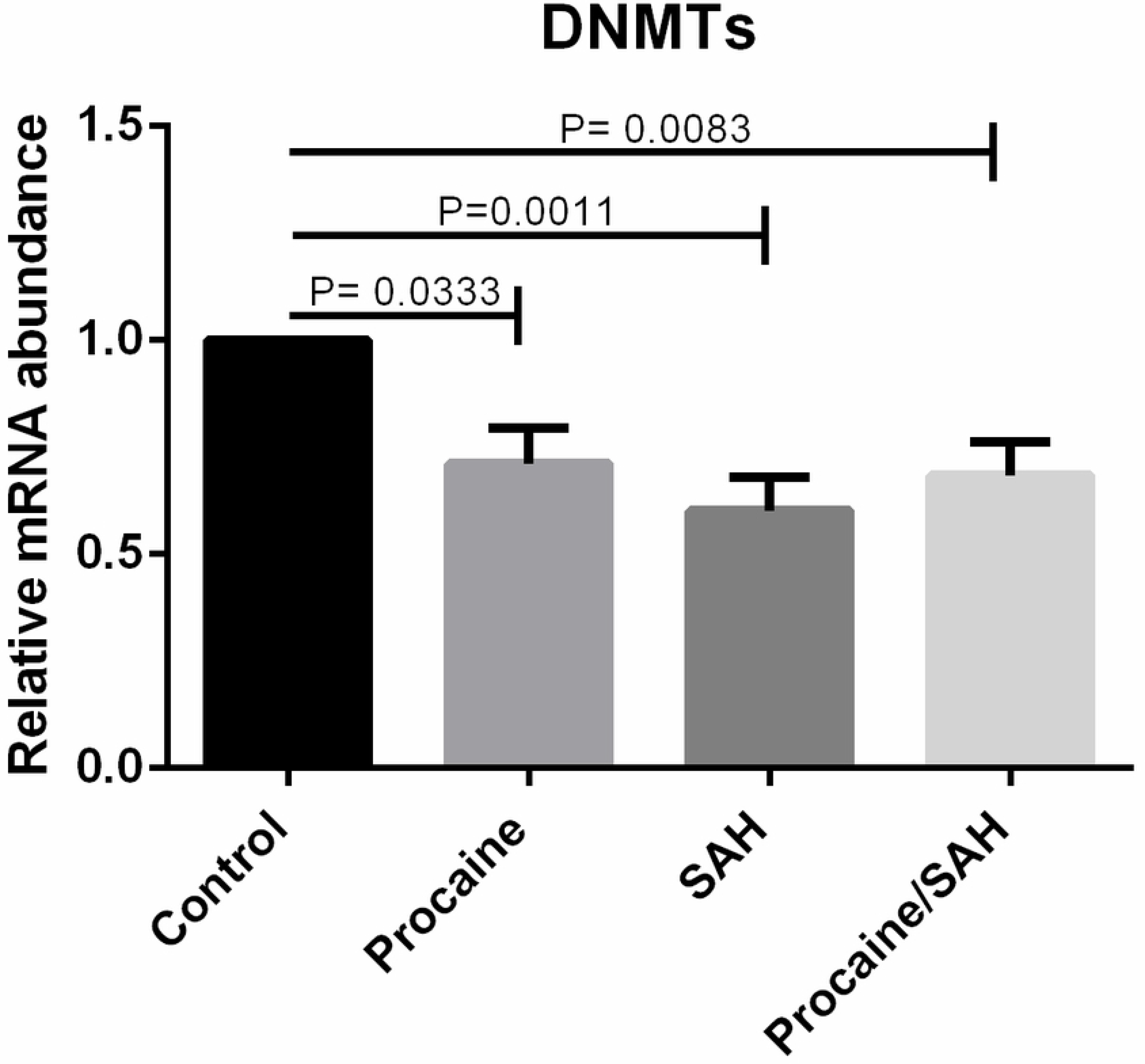
mRNA levels of the group of DNMTs (DNMT1, DNMT3A, DNMT3B) determined by RT-qPCR in bovine skin fibroblasts treated with procaine and/or S-adenosyl L-homocysteine (SAH). Differences were considered significant when p <0.05.

**Fig. 11.**
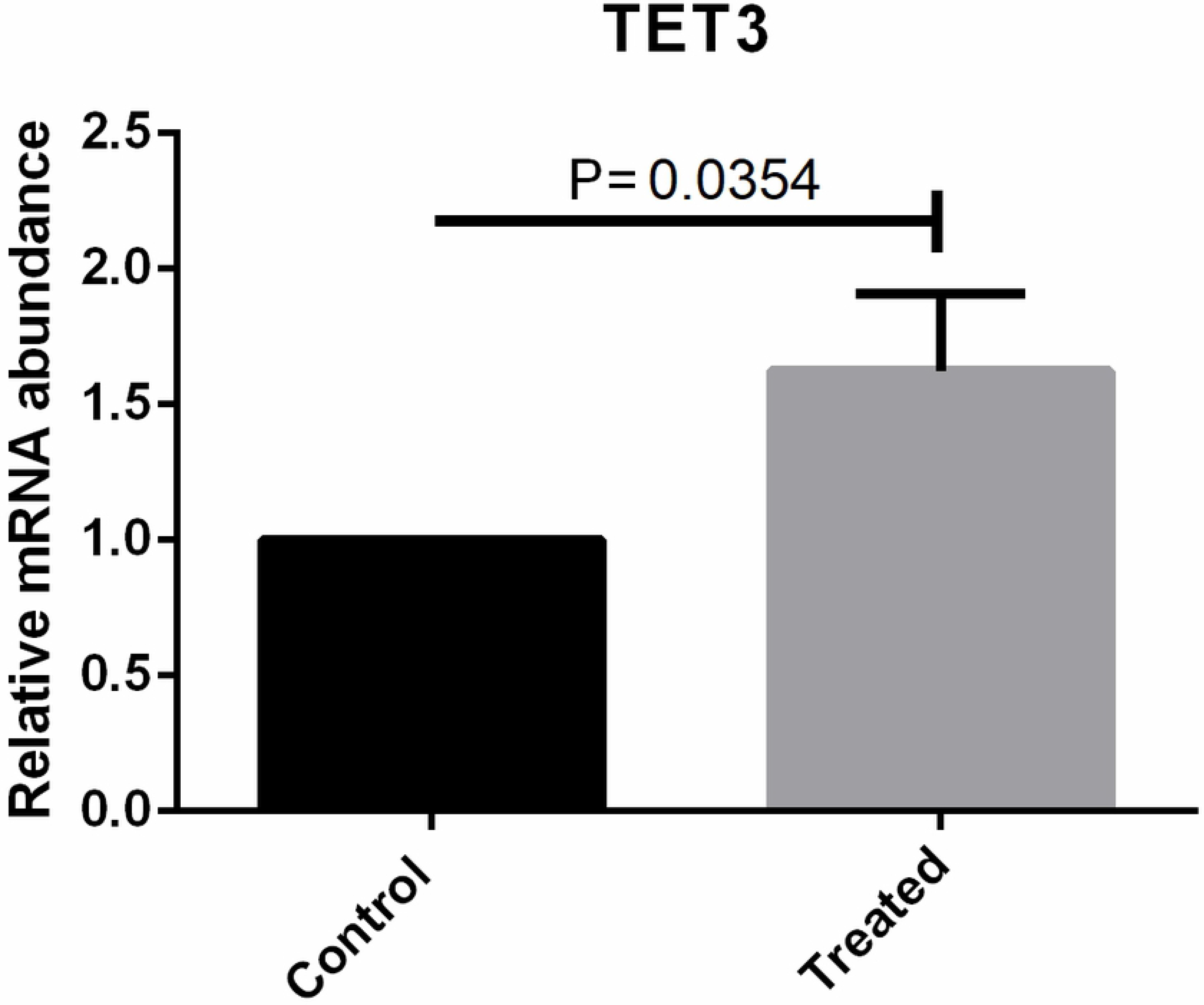
mRNA levels for the TET3 gene determined by RT-qPCR in bovine skin fibroblasts treated with procaine and/or S-adenosyl L-homocysteine (SAH). Differences were considered significant when p <0.05.

**Fig. 12.**
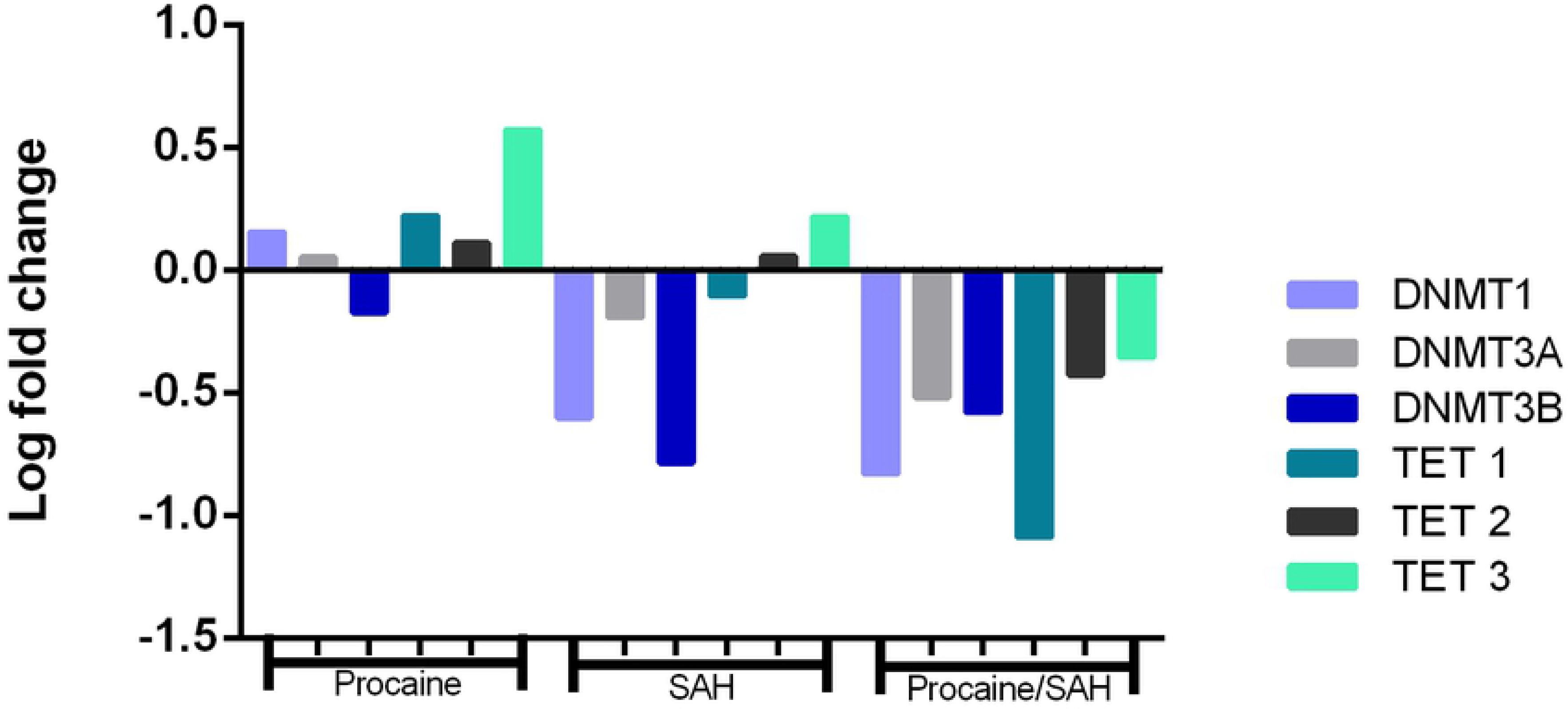
Fold change values for the DNMT1, DNMT3A, DNMT3B, TET1, TET2, and TET3 genes in skin fibroblasts cultured *in vitro* for 14 d with procaine, S-adenosyl L-homocysteine (SAH), or both substances in relation to the control group.

**Fig. 13.**
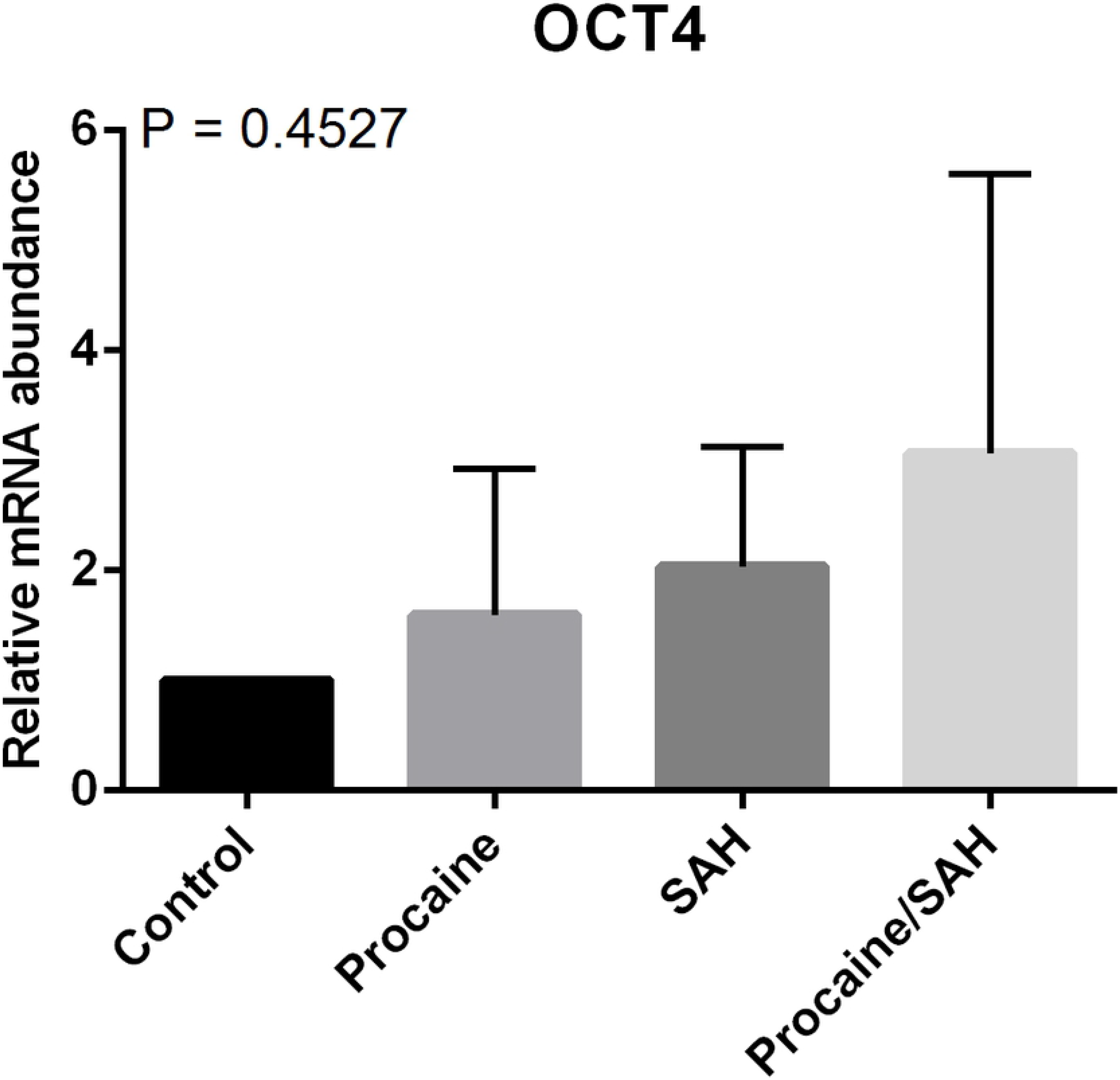
mRNA levels for the OCT4 gene determined by RT-qPCR in bovine skin fibroblasts treated with procaine and/or S-adenosyl L-homocysteine (SAH). Differences were considered significant when p <0.05.

**Fig. 14.**
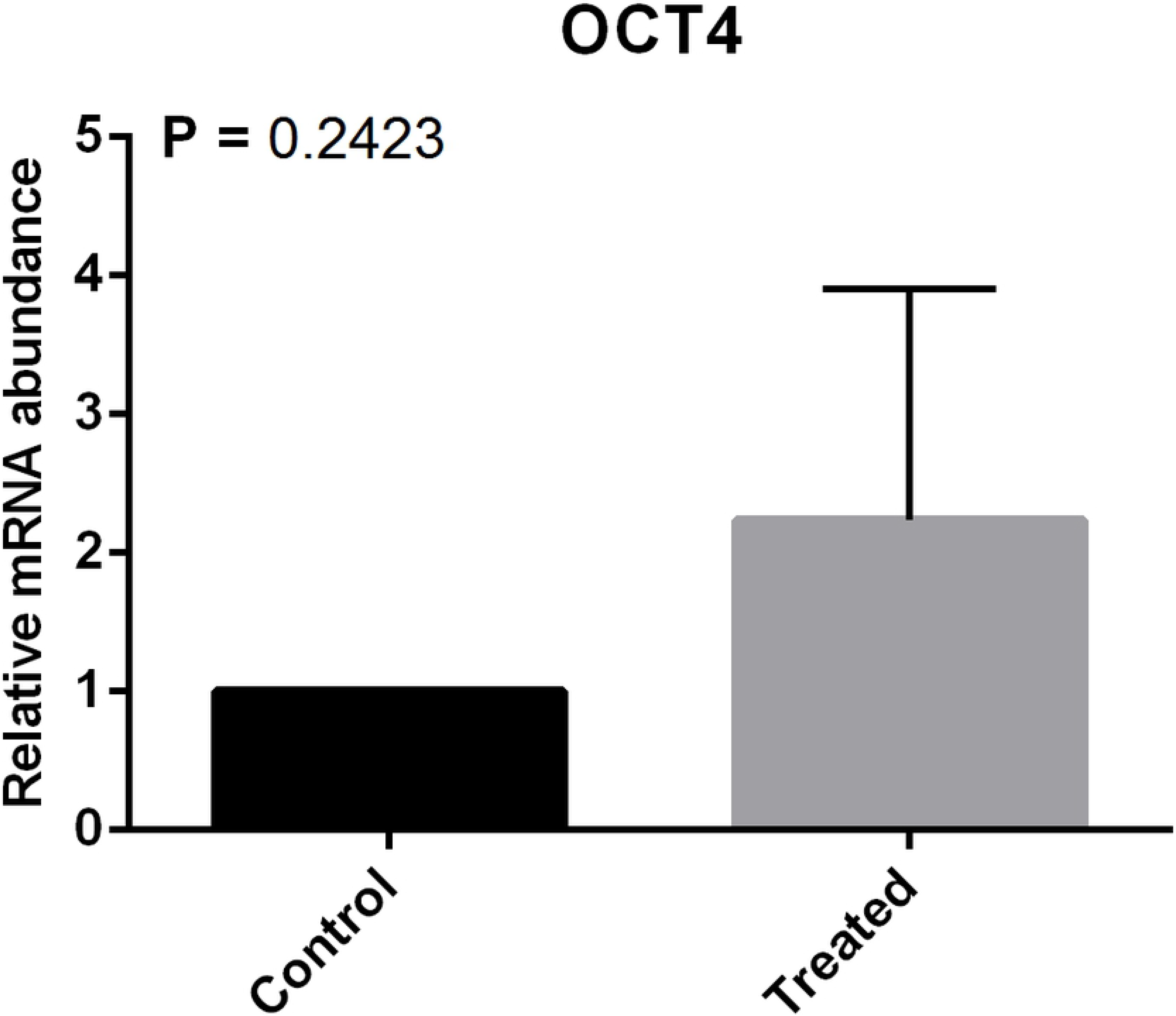
mRNA levels for the OCT4 determined by RT-qPCR in bovine skin fibroblasts treated with procaine and/or S-adenosyl L-homocysteine (SAH) regardless the treatments. Differences were considered significant when p <0.05.

## Discussion

Widespread epigenetic reprogramming, including changes in DNA methylation and post-translational histone modifications, occurs during mammalian gametogenesis and initial embryogenesis (9, 10). Most evidence suggests that the developmental failures and abnormalities in SCNT embryos and foetuses may be due to the incomplete epigenetic reprogramming of the somatic genome from the donor cell (7). Similar to the somatic genome of the donor cells, the genomes of SCNT embryos are hypermethylated compared to those of embryos produced *in vitro* or *in vivo* (5, 7). Thus, a strategy that decreases DNA methylation levels in somatic cells in culture could be useful for increasing SCNT efficiency.

Our results showed that the addition of procaine, SAH, or both compounds in combination to cell culture media did not impair cell growth or morphology (Fig. 2). Chromatin modulating agents that are analogous to nucleosides normally have high toxicity and cause apoptosis (23). Procaine and SAH are non-cytotoxic compounds that are able to inhibit DNMT enzymes (23, 34). Procaine was shown to reduce tumour volumes by 42.2%, not increase apoptosis rates, and have no adverse effects when used in the culture of the HLE hepatoblastoma cell line (35). Jeon et al. reported that SAH can reversibly inhibit DNMTs in somatic cells (24). Jafari et al. also reported that SAH induced global DNA demethylation (36). Taken together, these results support the potential use of procaine and SAH as demethylating agents in cell culture for different applications. Studies have shown that DNA methylation is an important epigenetic mark that plays a role in repressing gene activity, especially because it is associated with the chromatin state (37–39). Satellite DNA, such as satellite I, LINE-1, and α-satellite DNA, are normally methylated (40). Here, we showed that the DNA methylation of satellite I was lower in cells cultured with SAH (62.8%) and with SAH + procaine (64.2%) compared to that in control cells (72.5%) (Fig. 5). Thus, we believe that SAH may work well as a global DNA demethylating agent, especially considering that satellite I is widespread in the genome. We also found that SAH had a noticeable effect, at least in this repetitive region evaluated here, as it caused a significant reduction in methylation levels compared to those in the control, different from procaine, which resulted in levels similar to those in the control (Fig. 5). This higher level of methylation observed in the control group in the satellite region may be explained by the necessity for transcriptional repression by the methylation marks on the repetitive DNA, which is in agreement with the literature (41). Interestingly, the CpG site at position 17 was totally demethylated in all sequenced clones in all treatment groups, suggesting that this pattern may have some biological role that was not investigated in this study (Fig. 4).

The global methylation analysis revealed that cells from all treatments were less methylated than the control cells (Fig. 6). Cells from the control group showed 2.9% global methylation. A very relevant reduction was observed in the cells treated with procaine (1.3%), SAH (1.6%), and both compounds in combination (1.5%), demonstrating the significant potential demethylating action of both substances on bovine fibroblasts. We also found no differences among the groups that were treated with procaine, SAH, or both compounds.

The presence of tissue-specific epigenetic marks in somatic/differentiated genomes renders essential genes related to embryo development transcriptionally inactive, leading to abnormal global gene expression patterns in embryos (7, 37). Moreover, the incomplete DNA methylation reprogramming normally observed in SCNT could be caused by repressive factors present in donor cells (17, 42). Therefore, our proposal to reduce the methylation levels of the donor cell genome could improve reprogramming to more effectively erase cellular memory and promote the better activation of developmental genes.

Studies evaluating the expression profiles of genes encoding the enzymes related to the DNA methylation process in bovine fibroblasts are scarce. However, knowledge of the gene expression profile in somatic cells commonly used for the donor nucleus in SCNT may be important for efforts to improve the SCNT efficiency. The epigenomic state of early-developing human embryos is defined and governed by a group of genes that encode the enzymes involved in chromatin remodelling (41). Min et al. (2015) observed that the expression of genes involved in initial development were more altered in blastocysts produced using fibroblasts for the donor nucleus than in embryos obtained from cumulus cells (43). In addition, they also found that cloned embryos showed an altered gene expression profile compared to embryos produced *in vitro* (43).

In this study, we also evaluated the expression profile of genes related to DNA methylation reprogramming. The log-fold change analysis revealed that the overall profile of the genes encoding the DNMTs was downregulated in cells treated with SAH or a combination of procaine and SAH (Fig. 12). In addition, when we compared the mRNA levels of the group of three DNMT enzymes or compared the levels of transcripts between treated cells and control cells, we observed that the treatments decreased the DNMT mRNA levels (Fig. 9 and Fig. 10). At the cellular level, the enzymatic concentration may influence its own transcription rate. Molecules that are able to interact with enzymes in their active form may influence their activity (44, 45). Thus, this may lead to the initial accumulation of these enzymes, inducing the cell to then reduce the transcription rate (45). This mechanism could explain our results, in which we found a reduction in mRNA levels for DNMTs in the presence of procaine and SAH.

In addition to DNMTs, we also evaluated the levels of gene transcripts encoding TET enzymes; these act in the DNA demethylation process, and their correct expression is essential for embryo development (14, 46). Although it has been discussed in the literature in different contexts (47–50), reports on the expression profile of TET genes in cattle are scarce, especially those focusing on cloning using NT. TET enzymes oxidise 5-mC to 5-hmC, and in general, 5-hmC is transient in the epigenome, as its frequency is less than that of 5-mC (51). However, some cells have high levels of TETs, such as embryonic stem cells (52) and neurons (53). Here, we found higher levels of TET3 transcripts in cells cultivated with both substances, individually or together, compared to those in the control (Fig. 11). Studies with bovine embryos have shown that the TET1 enzyme does not participate in the DNA demethylation process in early development, with only TET2 and TET3 being detected and TET3 being present at higher levels in embryos (54) and oocytes (55). Our results demonstrated that TET3 expression increased in cells treated with SAH and procaine. Pagage-Lariviere and Sirard (2014) showed that TET3 is present in high abundance in embryos (54). In this context, the increased levels of TET3 found here may be favourable for the development of cloned embryos when using these cells as a nucleus donor.

Interestingly, we found that fibroblasts from all treatments expressed OCT4, even fibroblasts from the control group, even though there were not expected to express genes related to pluripotency because of their highly differentiated status. This result may be explained by the results of Pan et al. (2015) from their evaluation of skin fibroblast samples showing the presence of a heterogeneous population of fibroblasts containing multipotent stem cells (56). Moreover, we found no significant differences in OCT4 transcript levels among the treatments, suggesting that the molecular changes induced by procaine and SAH were not sufficient to alter the expression of pluripotency genes such OCT4. Because the control of gene expression is a mechanism involving many factors such as DNA methylation, histone modifications, and microRNAs, just the DNA demethylation induced by procaine and SAH might not have been sufficient to alter OCT4 expression.

In general, our data suggest that the use of procaine, SAH, or both compounds in combination is a promising strategy for SCNT protocols because their use in cell culture did not affect cell growth and morphology and because they reduced specific and global DNA methylation levels. However, it is important to highlight that the reduction in global methylation was higher than that for the methylation of the satellite region. This result may be explained by the fact that these repetitive regions are more resistant to being demethylated (57). A recent study demonstrated that Dnmt1s, a specific isoform in somatic cells, is a barrier for zygotic genome activation and genomic methylation reprogramming, leading to a developmental stop in SCNT embryos (18). Moreover, its removal allows for an efficient activation of genes essential for embryonic development, thus increasing SCNT efficiency (18). Based on this information, the use of procaine and SAH can be a useful strategy to improve SCNT efficiency considering that our results showed that these compounds decreased DNMT transcript levels (Fig. 9). Taken together, the use of these DNA demethylating agents in cell culture decreased DNMT expression and DNA methylation levels, suggesting that these agents have the potential to induce cells into a less differentiated state, which may improve SCNT protocols.

## Authors’ contributions

NAB participated in the collection of materials for analysis. NAB, AdosSM, MMS, LNV, LOL, and RVS performed the cellular and molecular analyses. MMF designed the study. NAB and MMF interpreted the results and wrote the manuscript. All authors read and approved the final manuscript.

## Acknowledgments

We would like to thank Capes, Brazil; Embrapa Genetic Resources and Biotechnology, Brazil; and FAP-DF for the support provided for this study.

